# Predicting future cognitive decline from non-brain and multimodal brain imaging data in healthy and pathological aging

**DOI:** 10.1101/2020.06.10.142174

**Authors:** Franziskus Liem, Kamalaker Dadi, Denis A. Engemann, Alexandre Gramfort, Pierre Bellec, R. Cameron Craddock, Jessica S. Damoiseaux, Christopher J. Steele, Tal Yarkoni, Daniel S. Margulies, Gaël Varoquaux

## Abstract

Cognitive decline occurs in healthy and pathological aging, and both may be preceded by subtle changes in the brain — offering a basis for cognitive predictions. Previous work has largely focused on predicting a diagnostic label from structural brain imaging. Our study broadens the scope of applications to cognitive decline in healthy aging by predicting future decline as a continuous trajectory, rather than a diagnostic label. Furthermore, since brain structure as well as function changes in aging, it is reasonable to expect predictive gains when using multiple brain imaging modalities. Here, we tested whether baseline multimodal neuroimaging data improve the prediction of future cognitive decline in healthy and pathological aging. Non-brain data (including demographics and clinical and neuropsychological scores) were combined with structural and functional connectivity MRI data from the OASIS-3 project (N = 662; age = 46 – 96y). The combined input data was entered into cross-validated multi-target random forest models to predict future cognitive decline (measured by the Clinical Dementia Rating and the Mini-Mental State Examination), on average 5.8y into the future. The analysis was preregistered and all analysis code is publicly available. We found that combining non-brain with structural data improved the continuous prediction of future cognitive decline (best test-set performance: R^2^ = 0.42) and that cognitive performance, daily functioning, and subcortical volume drove the performance of our model. In contrast, including functional connectivity did not improve predictive accuracy. In the future, the prognosis of age-related cognitive decline may enable earlier and more effective cognitive, pharmacological, and behavioral interventions to be tailored to the individual.

## 1 Introduction

Cognitive decline, such as worsening memory or executive functioning, occurs in healthy and pathological aging. Crucially, noticeable decline may be preceded by subtle changes in the brain. It is this sequence that enables using brain imaging data to predict the current cognitive functioning of a person or related surrogate markers. For example, structural brain imaging has been used to predict patients’ current cognitive diagnosis (Rathore et al., 2017), or brain-age (Cole and Franke, 2017), a surrogate biomarker related to cognitive impairment (Liem et al., 2017). Together, these findings demonstrate the clinical potential of neuroimaging data used in combination with predictive analyses.

While predicting *current* cognitive functioning enables insight into related brain markers, predicting *future* cognitive decline from baseline data poses a greater challenge with more substantial clinical relevance (Davatzikos, 2019). Using current brain imaging data to predict a current diagnostic label (such as dementia), targets a label that can fairly easily be determined via other means such as clinical assessments (and usually with less cost than brain imaging). When predicting future cognitive change, however, brain imaging might aid a prognosis with greater clinical utility that cannot be easily obtained otherwise. Most previous studies that predicted future change restricted their analysis to whether patients with mild cognitive impairment (MCI) converted to Alzheimer’s disease (AD) (Davatzikos et al., 2011; e.g., Eskildsen et al., 2015; Gaser et al., 2013; Korolev et al., 2016) or predicted membership in data-driven trajectory-groups of future decline (Bhagwat et al., 2018). Predicting future cognitive decline on a continuum (instead of forming distinct diagnostic labels from cognitive data) better characterizes the underlying change in abilities on an individual level. This approach can also be used to widen the scope of applications by including healthy aging. Brain data is a rich source of information that might help us better understand and even reorganize diagnostic syndromes or categories.

While most previous predictive studies used structural brain imaging alone, integrating structural and functional imaging has been shown to improve predictions. Since both brain structure (Oschwald et al., 2019) and brain function (Liem et al., 2020) change in aging, the most accurate predictions of brain-age have come from combining them (Engemann et al., 2020; Liem et al., 2017). Multimodal gains have also been shown in more complex predictions such as current diagnosis in AD (Rahim et al., 2016) and conversion from MCI to AD (Dansereau et al., 2017; e.g., Hojjati et al., 2018). Therefore, integrating multiple brain imaging modalities enables a more complete characterization of bran aging and provides increased predictive power.

The present study aimed to predict future cognitive decline from baseline data in healthy and pathological aging. We combined non-brain data, such as scores from clinical assessments and demographics, with multimodal brain imaging data to test whether adding brain imaging to non-brain data improves predictive performance, and whether multimodal imaging outperforms single imaging modalities. We showed that structural imaging in particular improved continuous prediction of future cognitive decline. An early prognosis of future cognitive decline might enable earlier and more effective pharmacological or behavioral treatments to be tailored to the individual, resulting in more efficiently allocated medical resources.

## 2 Methods

The analysis presented here was preregistered (Liem et al., 2019). We largely followed this preregistration and deviations are described in the supplement (*6.1.2 Deviation from preregistration*). The deviations concern minor details in data analysis and do not affect the qualitative conclusions we draw. Additionally, we performed non-preregistered validation analyses that were suggested by the main results.

### 2.1 Sample and session selection

The present analysis aimed to predict future cognitive decline from baseline non-brain (e.g., age and clinical scores) and brain imaging data (regional brain volume and functional connectivity). We used data from the publicly available, longitudinal OASIS-3 project, a collection of data from several studies at the Washington University Knight Alzheimer Disease Research Center (LaMontagne et al., 2019). OASIS-3 acquired data in different types of sessions (*clinical sessions:* non-brain data describing personal characteristics, cognitive and everyday functioning, health; *neuropsychological sessions:* non-brain data from neuropsychological tests; *MRI sessions:* structural and functional MRI). The count and spacing between sessions varied between subjects. To predict future cognitive decline, baseline sessions were used as input data and follow-up sessions as targets. The study design required a matching approach to select i) baseline sessions (from clinical, neuropsychological, and MRI sessions) to be used as input data, and ii) follow-up clinical sessions to estimate the future cognitive decline.

First, baseline data were established by matching sessions from the different types (clinical, neuropsychological, MRI). We matched each MRI session that had at least one T1w and one fMRI scan with the closest clinical session. For each subject, the first MRI-clinical-session pair with an absolute time difference < 1 year was selected as baseline session. If no such pair was available, the subject was excluded from the analysis. Additionally, the closest neuropsychological session (within 1 year of the MRI baseline session) was also considered as baseline data. Baseline information from neuropsychological testing, however, was considered optional and not finding a matching neuropsychological session was not a criterion for exclusion. All data preceding the selected baseline sessions were disregarded for the analysis.

Second, all clinical sessions after the baseline clinical session were included as follow-up sessions to estimate cognitive decline. To reliably estimate decline, subjects were only included if they had at least three clinical sessions (baseline plus two follow-up sessions).

This matching approach reduced the sample (N_total_=1098) to 662 subjects (302 male; Table 1)^1^. The majority was cognitively healthy at baseline (509 healthy controls, 12 were diagnosed with MCI, and 111 with dementia; for 30 no diagnosis was available for the baseline session).

**Table 1.** Sample characteristics. N = 662 (302 male). MMSE: Mini-Mental State Examination

|  | M | SD | min | max |
| --- | --- | --- | --- | --- |
| age<sub>baseline</sub> | 71 | 8 | 46 | 96 |
| MMSE<sub>baseline</sub> | 28 | 2 | 16 | 30 |
| N clinical sessions | 5.8 | 2.4 | 3 | 15 |
| years between clinical sessions | 1.2 | 0.6 | 0.003 | 5.5 |
| years in study | 5.8 | 2.5 | 1.6 | 10.9 |

MRI data was downloaded in BIDS format (Gorgolewski et al., 2016) via scripts provided by the OASIS project^2^. Non-brain data was downloaded via XNAT central^3^.

### 2.2 Data

#### 2.2.1 Non-brain data

Non-brain data described personal characteristics at baseline, such as demographics, cognitive and everyday functioning, genetics, and health (Table 2 shows the abbreviations of tests). For further information on the measurements, see relevant publications by the OASIS team (LaMontagne et al., 2019; Morris et al., 2006; Weintraub et al., 2009).

**Table 2.** List of abbreviations of clinical tests

|  |  |
| --- | --- |
| CDR | Clinical Dementia Rating |
| FAQ | Functional Activities Questionnaire |
| GDS | Geriatric Depression Scale |
| MMSE | Mini-Mental State Examination |
| NPI-Q | Neuropsychiatric Inventory Questionnaire |
| TMT | Trail Making Test |
| WAIS-R | Wechsler Adult Intelligence Scale-Revised |
| WMS-R | Wechsler Memory Scale-Revised |
| BNT | Boston Naming Test |

The specific measures included:

1. demographic information: sex, age, education
2. clinical scores: MMSE (Folstein et al., 1975), CDR (Morris, 1993), FAQ (Jette et al., 1986), NPI-Q (Kaufer et al., 2000), GDS (Geriatric Depression Scale, Yesavage et al., 1982)
3. neuropsychological scores: WMS-R (Elwood, 1991), Word fluency, TMT (Heller et al., 2013), WAIS-R (Franzen, 2000), BNT (Borod et al., 1980)
4. APOE genotype
5. a cognitive diagnosis (healthy control, MCI, dementia)
6. health information: cardio/cerebro-vascular health, diabetes, hypercholesterolemia, smoking, family history of dementia
7. the number of clinical session conducted before the selected baseline session (for instance sessions without a matching MRI session) to account for retest effects

#### 2.2.2 MRI data

MRI data were acquired on Siemens 3T scanners, with the majority coming from a TrioTrim model (622 of 662 subjects). Each participant had between 1 and 4 T1w scans (1.7 on average). In total, the sample had 1’119 T1w images. The parameter combination most commonly used (in over 1’070 scans) was voxel size = 1 × 1 × 1 mm^3^, echo time (TE) = 0.003 s, repetition time (TR) = 2.4 s. Where available, T2w images were also used to aid surface reconstruction. In total, 618 participants had a T2w image. The parameters for the T2w images were voxel size = 1 × 1 × 1 mm^3^, TE = 0.455 s, TR = 3.2 s.

Each participant had between 1 and 4 functional resting-state scans (M = 2.0). In total, the sample had 1’327 functional images. The parameter combination most commonly used (in over 1’300 scans) was voxel size = 4 × 4 × 4 mm^3^, TE = 0.027 s, TR = 2.2 s, scan duration = 6 min. For further information regarding the imaging data see (LaMontagne et al., 2019).

### 2.3 MRI preprocessing

Functional and structural MRI data were preprocessed using the standard processing pipeline of *fMRIPrep* 1.4.1 (Esteban et al., 2018b), which also includes running *FreeSurfer* 6.0.1 on the structural images (Fischl, 2012). A detailed description of the preprocessing can be found in the supplement (*6.1.1 Details on MRI preprocessing*). Except for basic validity checks in a random subset of subjects, data quality of the preprocessed data was not rigorously assessed. Notably *fMRIPrep* has been shown to robustly work across many datasets (Esteban et al., 2018b).

### 2.4 Feature extraction

Input data from non-brain and brain imaging modalities at baseline were used to predict future cognitive decline (predictive targets). In the following sections we provide further details on the features that entered the predictive models.

#### 2.4.1 Input data

Input data for the predictive models came from three modalities: *non-brain*, global and subcortical structural (*structGS*), and functional connectivity (*func*; Figure 1-1). Modalities were entered into the models on their own and in combination. For instance, *non-brain + structGS* models received horizontally concatenated input features from the *non-brain* and *structGS* modalities (Figure 1-2). This allowed testing whether combining non-brain with structural data improved predictive accuracy as compared to non-brain data alone. The following paragraphs describe the input data modalities and Table S1 gives an overview of features entered into the models.

**Figure 1.**
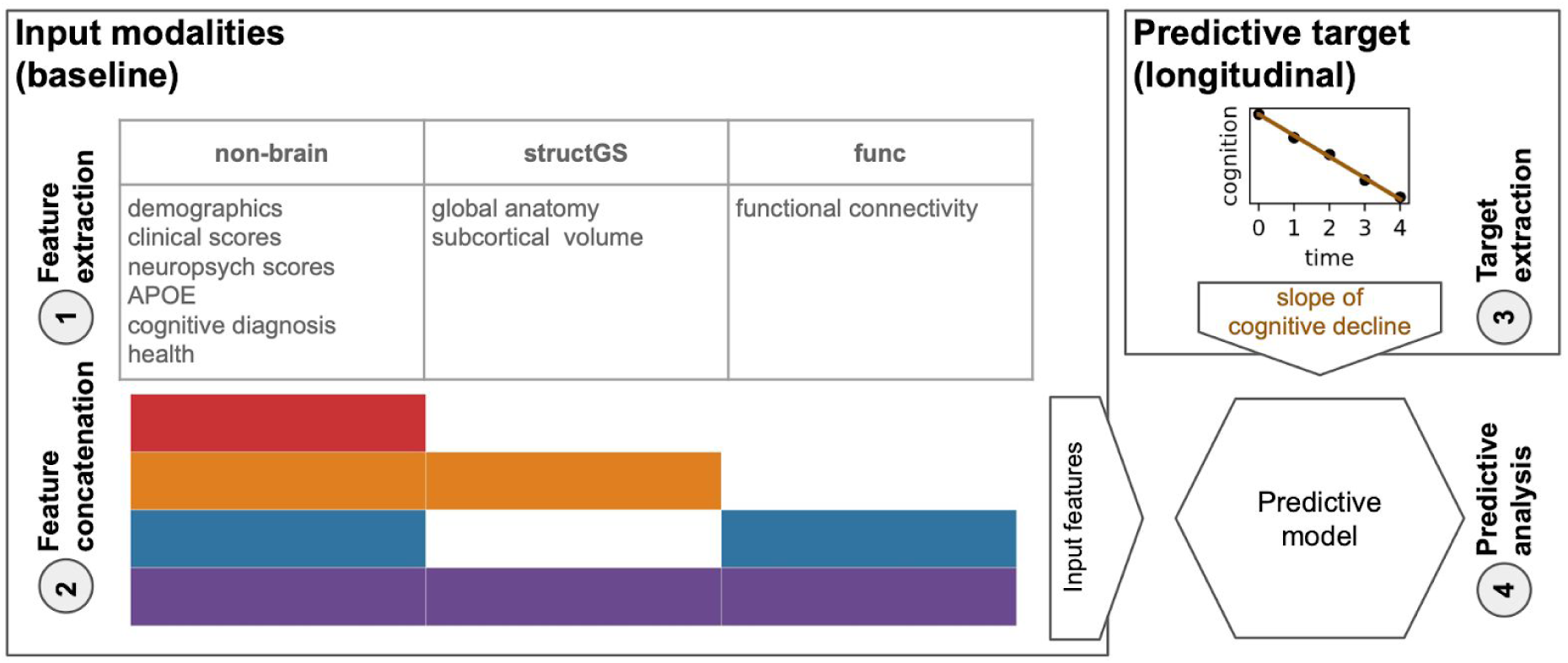
Overview of the predictive approach. 1) Features from *non-brain, structGS* (global and subcortical structural), and *func* (functional connectivity) modalities are extracted from baseline data. 2) Feature concatenation produces sets of multimodal input features. For instance, red represents *non-brain* features only, while orange represents a combination of *non-brain* and *structGS*. 3) Extraction of slopes representing cognitive change from CDR (Clinical Dementia Rating) and MMSE (Mini-Mental State Examination). 4) Models are trained to predict the cognitive decline based on the input features. Here, we used a multi-target random forest model within a nested cross-validation approach to predict CDR and MMSE change simultaneously.

**Figure 2.**
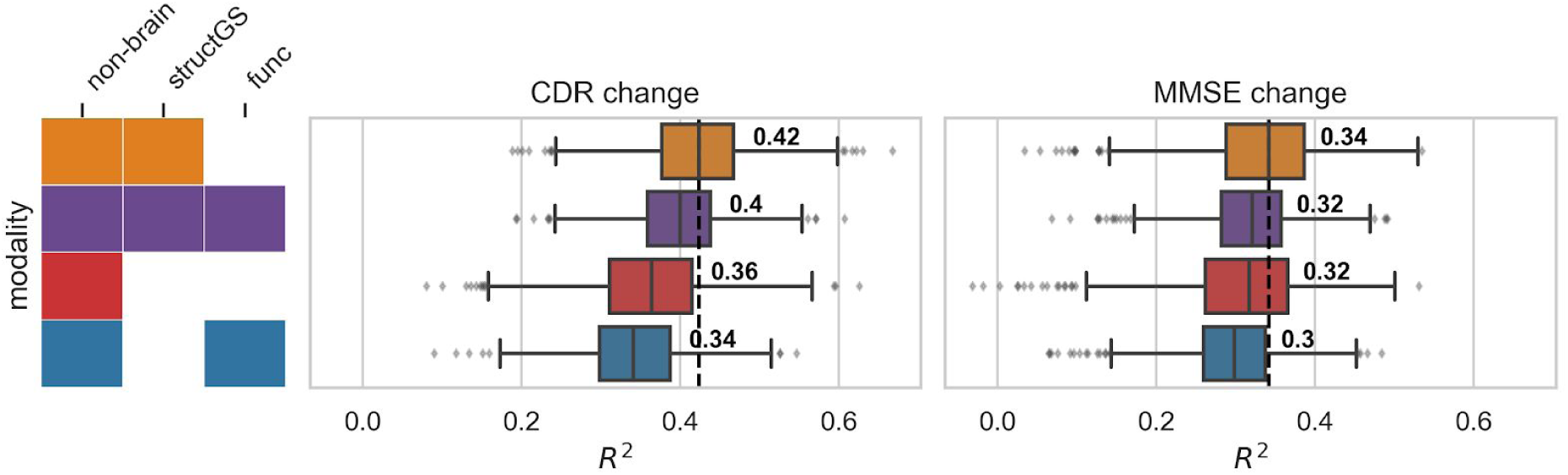
Adding structural data (orange) to non-brain data (red) improved the prediction of cognitive decline. Test performance (R^2^, coefficient of determination, x-axis) across splits (N_splits_ = 1000) for the combinations of input modalities (y-axis). Targets: cognitive change measured via CDR (Clinical Dementia Rating, middle) and MMSE (Mini-Mental State Examination, right). Input modalities: non-brain, structGS (global and subcortical structural volumes), func (functional connectivity). The left panel represents combinations of input modalities (e.g., orange is *non-brain + structGS*). The number represents the median, the dashed vertical line marks the median of the best-performing combination of modalities (within a target measure). For the full results that include single-modality brain imaging, see Figure S2.

##### 2.4.1.1 Non-brain data

Non-brain features included demographics, scores of clinical and neuropsychological instruments, APOE genotype, and health information. For a detailed list see Table S1. In total, 66 features entered the models from the *non-brain* modality.

##### 2.4.1.2 Structural MRI (structGS)

For the *structGS* modality (global and subcortical structure), anatomical markers were extracted from the *FreeSurfer*-preprocessed anatomical scans. Following our previous work (Liem et al., 2017), we extracted global structural markers (volume of cerebellar and cerebral GM and WM, subcortical GM, ventricles, corpus callosum, and mean cortical thickness) and the volumes of seven subcortical regions (accumbens, amygdala, caudate, hippocampus, pallidum, putamen, thalamus; for each hemisphere separately). Most markers were extracted from the *aseg* file, except for mean cortical thickness, which was extracted from the *aparc.a2009s* parcellation (Desikan et al., 2006). To account for head-size-effects, volumetric values were normalized by estimated total intracranial volume. In total, 35 features entered the models from the *structGS* modality.

##### 2.4.1.3 Functional MRI (func)

Functional connectivity was computed from the *fMRIPrep*-preprocessed functional scans. Denoising was performed using the 36P model (Ciric et al., 2017), which includes signals from 6 motion parameters, global, white matter, and CSF signals, derivatives, quadratic terms, and squared derivatives. Time series were extracted from 300 cortical, cerebellar, and subcortical coordinates of the Seitzman atlas (Seitzman et al., 2018) using balls of 5 mm radius. The signals were band-pass filtered (0.01-0.1 Hz) and linearly detrended. Connectivity matrices were extracted by correlating the time series using Pearson correlation and applying Fisher-z-transformation. If multiple fMRI runs were available, the z-transformed connectivity matrices were averaged within subjects. The vectorized upper triangle of this connectivity matrix was entered into the predictive pipeline and was further downsampled to 100 PCA components within cross-validation (see below). Denoising and feature extraction was performed with *Nilearn* 0.6.0 (Abraham et al., 2014).

#### 2.4.2 Predictive targets

To quantify future cognitive decline, trajectories of two clinical assessments, the CDR (Clinical Dementia Rating, Sum of Boxes score) and the MMSE (sum score of the Mini-Mental State Examination) were estimated using an ordinary least squares linear regression model for each subject and assessment independently (Figure 1-3; for information on the count and timing of sessions, see Table 1). A linear slope was fitted through the raw scores of the follow-up session with the intercept fixed at the raw score of the baseline session (*score*_*fu, a*_ ∼ *score*_*bl, a*_ + β_*slope, a*_ × *time*; fu: follow-up, a: assessment, bl: baseline). This approach was chosen over a linear mixed effects model, as the mixed effects model requires data from multiple subjects, making cross-validation more convoluted. The resulting two parameters (β_*slope, CDR*_ and β_*slope, MMSE*_) were the two targets that were simultaneously predicted in the predictive analysis using a multi-target approach (Rahim et al., 2017). Slopes were estimated with *Statsmodels* 0.10.1 (Seabold and Perktold, 2010). The distribution of the estimated targets is plotted in Figure S1.

**Figure 3.**
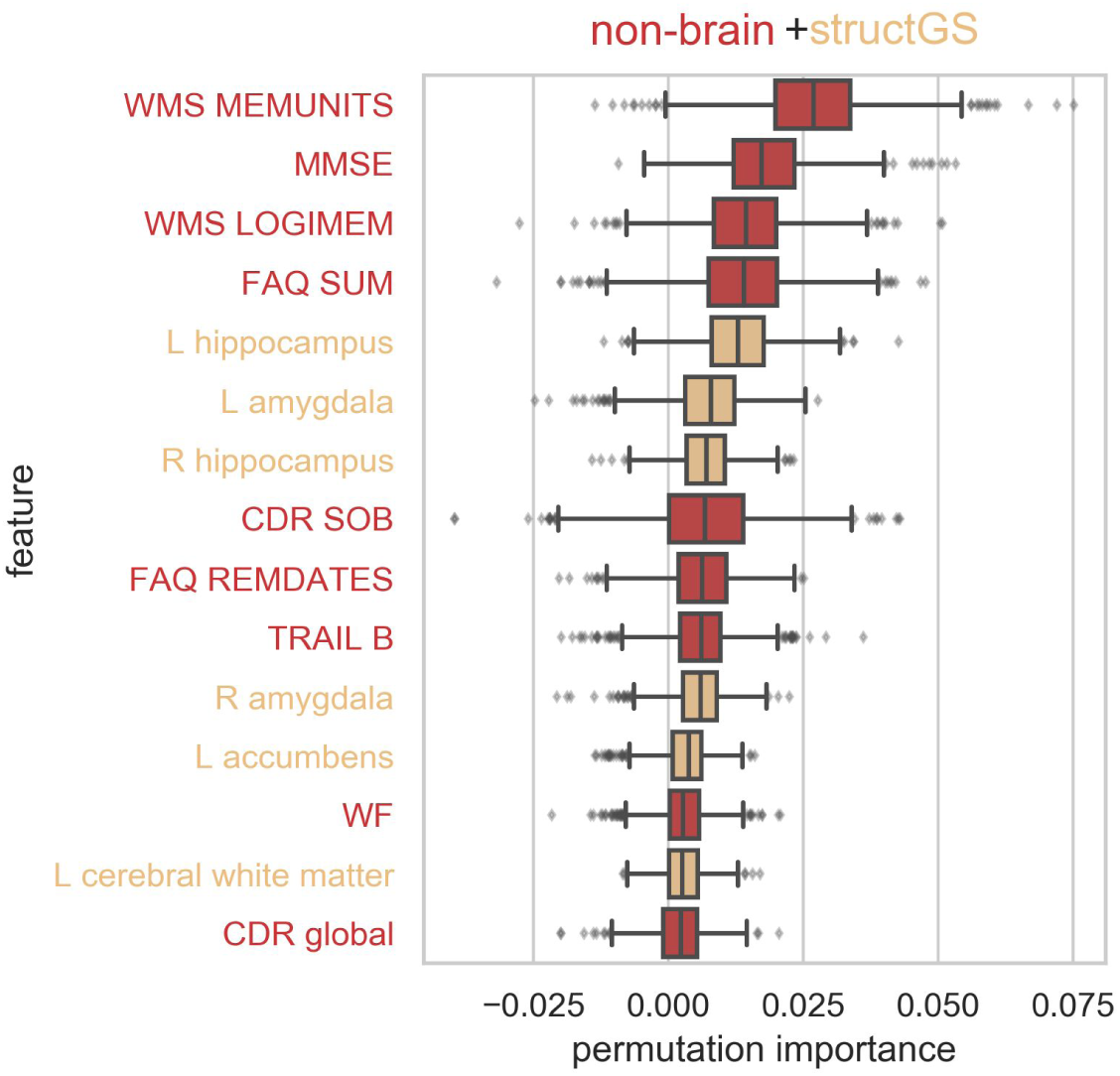
Cognitive performance, daily functioning, and subcortical volume were among the most informative features. Permutation importance of the top 15 features of the *non-brain + structGS* model (median across splits). Permutation importance is quantified as the decrease in test performance R^2^ with the feature permuted. Red: non-brain features, light orange: structGS features. CDR: Clinical Dementia Rating, FAQ: Functional Assessment Questionnaire, L: left, MMSE: Mini-Mental State Examination, R: right, SOB: sum of boxes score, TRAIL B: Trail Making Test B, WF: word fluency, WMS: Wechsler Memory Scale.

### 2.5 Predictive analysis

The predictive pipeline (Figure 1-4) consisted of a multivariate imputer (*Scikit-learn’s* IterativeImputer) (Buck, 1960) and a multi-target random forest (RF) regression model (Breiman, 2001). Multivariate imputation has recently been shown to robustly work in combination with predictive models under different missingness scenarios (Josse et al., 2019).

Predictive models were trained using nested cross-validation via a stratified shuffle-split (1000 splits, 80% training, 20% test subjects, stratified by the targets). In the inner loop, the RF’s hyperparameters were tuned via grid search on the training subjects (the tree depth was selected among [3, 5, 7, 10, 15, 20, 40, 50, None], where None leads to fully grown trees; the criterion to measure the quality of an RF-split was tuned with ‘mean squared error’ and ‘mean absolute error’). The best estimator was carried forward to determine its out-of-sample performance on the test subjects. To derive an estimate of chance performance, null-models were also trained and tested with permuted target values. For each cross-validation split, the coefficient of determination (R^2^) was calculated on the test predictions. All predictive analyses were performed using *Scikit-learn* 0.22.1 (Pedregosa et al., 2011).

Model comparison was used to determine whether one model offered better prediction accuracy than another (for instance, to check whether a given model outperformed the permuted model, or whether a model with added brain imaging data improved accuracy as compared to a model using only non-brain data). Model comparison in a cross-validation needs to take the dependence between splits into account, complicating statistical tests (Bengio and Grandvalet, 2004). Thus, instead of calculating a formal statistical test, we calculated the number of splits for which the model in question outperformed the reference model, resulting in a percent value, with numbers close to 100% denoting models which robustly outperformed the reference model (Engemann et al., 2020).

To inspect which features contributed to a prediction, permutation importance was calculated (Breiman, 2001). Permutation importance evaluates the effects of features on the predictive performance by permuting feature values. If shuffling a feature does decrease performance, it is considered important for the model. It has to be noted that this approach might underestimate the importance of correlated features. Furthermore, learning curves were estimated to assess whether the number of subjects in the analysis was sufficient. For these comparisons, models were trained with increasing sample size while observing the test performance.

We performed additional analyses to diagnose the predictive pipeline and present our results in context. First, to validate the pipeline and analysis code, the same predictive methodology was used to predict age, a strong and well-established effect (Liem et al., 2017). For this validation, age was removed from the input data and the approach followed in the main analysis was repeated using a ridge regression model and 200 cross-validation splits.

Second, to better compare our results with previous work that predicted decline using class labels, we repeated the original pipeline to classify extreme groups of subjects that are cognitively stable vs. subjects with cognitive decline using random forest classifiers and 200 cross-validation splits. Subjects with CDR-SOB slopes > 0.25 were labeled as declining (N = 156), and a randomly drawn equal number of subjects without change in CDR-SOB were labeled as stable.

### 2.6 Open science statement

All data used in the analysis are publicly available via the OASIS-3 project (LaMontagne et al., 2019). The analysis plan was preregistered (Liem et al., 2019). All preprocessing and analyses were performed in *Python* using open-source software and the code for preprocessing and predictive analysis is publicly available (Liem, 2020)^4^. Furthermore, a docker container which includes all software and code to reproduce the preprocessing and predictive analysis is also provided^5^.

## 3 Results

### 3.1 Predicting cognitive decline

A combination of non-brain and structural data gave the best predictions of future cognitive decline. Adding structural data improved the prediction for both the CDR (median test performance R^2^ increased from 0.36 to 0.42; Figure 2, red vs. orange; for a scatter plot showing true vs predicted values, see Figure S3) and the MMSE (0.32 to 0.34), as compared to predictions from non-brain data alone. This increase occurred in a large majority of splits (91% of splits for CDR, 78% for MMSE; Table S2). In contrast, adding functional connectivity features to non-brain features, or to *non-brain + structGS* features, slightly decreased predictive performance.

To tune the RF models to the given problem, hyperparameters were optimized in a grid search approach. Tuning curves showed the results to be robust across a wide range of hyperparameter settings (Figure S4). Furthermore, learning curves demonstrated a sufficient sample size in the current setting (Figure S5).

The models consistently outperformed null models. Comparing the predictions against a null model with permuted predictions showed that most modalities outperformed chance-level in 100% of splits (Table S3). The predictions based on functional connectivity were an exception and outperformed null-models to a lesser degree (91% of splits for CDR, 73% for MMSE).

### 3.2 Features that predict cognitive decline

We used permutation importance to characterize the most predictive features of the best performing modality (non-brain + structGS). Within the top-15 features, non-brain included memory scores, the baseline scores of the targets (CDR, MMSE), and scores from the FAQ (functional assessment questionnaire). The structural features predominantly included subcortical regions (left and right hippocampus and amygdala, left accumbens; Figure 3).

### 3.3 Validation analyses

Although functional connectivity models predicted cognitive decline poorly, functional data improved accuracy when predicting brain-age. Since functional connectivity alone did not predict cognitive decline well and did not increase the predictive accuracy of the non-brain model (Figure S2), we conducted a validation analysis to ensure that our functional connectivity models were able to predict brain-age, a well-established surrogate biomarker. Here, we predicted age from the same input data as in the main analysis after first removing age from the input features set. In line with our expectations, functional connectivity increased predictive performance when combined with other modalities (e.g., in combination with non-brain, performance increased from 0.45 to 0.53; Figure 4), and functional connectivity alone could predict age reasonably well (median R^2^ = 0.33, Figure S6), suggesting that its negligible contribution to decline prediction cannot be attributed to general methodological or data quality issues.

**Figure 4.**
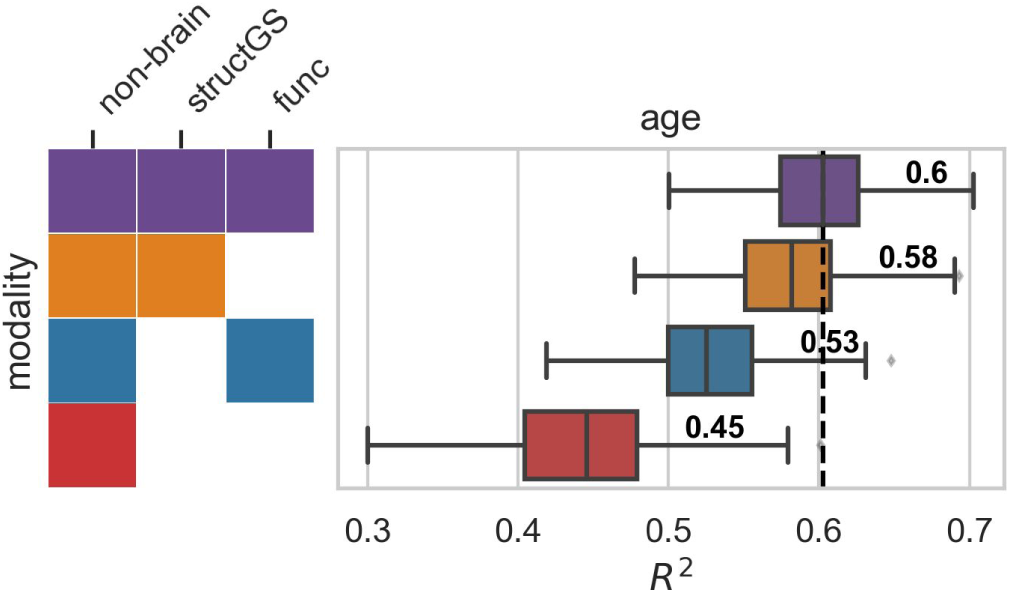
Multimodal imaging improves brain-age prediction. Input modalities: non-brain, structGS (global and subcortical structural volumes), func (functional connectivity). The number represents the median, the dashed vertical line marks the median of the best-performing combination of modalities. For the full results that include single-modality brain imaging, see Figure S6.

In the main analysis, the predictive target of cognitive decline was quantified as a continuous score. To compare our analysis to previous work that predicted classes of cognitive decline, we performed a further analysis that predicted extreme groups of cognitive decline (stable vs decline). Overall, extreme groups could be accurately predicted from the input data (most F1-scores [harmonic mean of the precision and recall] in the range of 0.8-0.9; Figure S7).

## 4 Discussion

In the present study, we found that combining baseline structural brain imaging data with non-brain data improved the prediction of future cognitive decline. In contrast, functional connectivity features did not improve prediction. By predicting future cognitive decline as a continuous trajectory, rather than a diagnostic label, our study broadens the scope of applications to cognitive decline in healthy aging. It also allows for more nuanced predictions on an individual level. In the future, these continuous measures may facilitate dimensional approaches to pathology (Cuthbert, 2014).

The benefit of combining structural with non-brain data found in the present study is well in line with previous work that predicted conversion from MCI to AD (Korolev et al., 2016), and classes of cognitive decline (Bhagwat et al., 2018). Non-brain data alone predicted cognitive decline and the model was robustly improved by adding structural data (R^2^ increased from 0.36 to 0.42 for CDR and from 0.32 to 0.34 for MMSE). These findings are consistent with prior work (Bhagwat et al., 2018; Korolev et al., 2016). In general, the range of accuracies reported in our study is well in line with previous work predicting a related continuous target (time to symptom onset in AD) (Vogel et al., 2018), as well as with work predicting diagnostic labels (Davatzikos et al., 2011; e.g., Eskildsen et al., 2015; Gaser et al., 2013; Korolev et al., 2016). After having established that a combination of non-brain and structural data gives predictions worthy of consideration, next, we assessed which features drove the predictions.

We found that clinical and neuropsychological assessments and subcortical structures drove the performance of our model. Measurements of memory, verbal fluency, executive function, and a wide set of cognitive and daily functions (MMSE, CDR, FAQ) were the most informative non-brain features for predicting cognitive decline. This matches well with Korolev et al. (2016) who found memory scores and clinical assessments (ADAS-Cog, FAQ) to be among the most informative non-brain features. On the other hand, hippocampus and amygdala volume were the most informative structural features in our analysis, which is well in line with previous work predicting conversion from MCI to AD (Eskildsen et al., 2015; Korolev et al., 2016). In contrast, risk factors (such as age, APOE, or health risks) and markers that quantify general brain atrophy and regional cortical brain structure did not add markedly to model performance. It should be noted that features were assessed using permutation importance, which underestimates the importance of correlated features. Alternative approaches, such as mean decrease impurity, might complement the permutation-based approach in future studies to improve the sensitivity (Engemann et al., 2020). Nevertheless, taken together, our results suggest that memory, everyday functioning, and subcortical features better predict future cognitive decline at the individual level than risk factors or global brain characteristics.

Functional connectivity, in contrast to brain structure, did not improve predictions when added to other modalities, nor did it predict cognitive decline on its own. While many previous studies predicted cognitive performance or decline based on structural imaging, studies using functional connectivity are rare and contain widely varying estimates of its predictive power (Dansereau et al., 2017; Hojjati et al., 2018; Vogel et al., 2018). Although functional connectivity in our study did not predict future cognitive decline, it did predict brain-age. Assuming that functional connectivity is at least somewhat predictive of future cognitive decline, our analysis may suggest that the processing of functional connectivity data was not a good fit for the cognitive targets. Furthermore, data with better spatial and temporal resolution might be able to better capture decline. This calls for future studies that benchmark different processing options as these can severely impact predictive accuracy (Dubois et al., 2018).

In the following, we will sketch possible future developments along four themes: implications of and possible improvements to the continuous *targets of cognitive decline, multimodal input data, predictive models*, and the importance of *generalization* to new datasets.

Quantifying cognitive decline continuously rather than discretely enables a more fine-grained and robust prediction, but also requires methodological choices. By predicting a diagnostic label, previous studies were often restricted to MCI patients and aimed to distinguish stable from converting patients. Considering decline as a continuum better characterizes the underlying change in abilities and allows for capturing changes that occur in healthy aging. Overcoming the scarcity of diagnosed conditions, this widens the scope of applications and has methodological advantages: the resulting increased sample size yields more robust models, which is critical to avoid optimistic bias in estimating prediction accuracy (Varoquaux, 2018; Woo et al., 2017). Furthermore, our approach also does not require assigning a diagnostic label, which entails subjective clinical judgment and arbitrary cut-off values. Considering cognitive decline as a continuous target does, however, require a model to aggregate multiple longitudinal measurements. Here, we used subject-specific linear slopes estimated through longitudinal data from clinical assessments. Since cognitive decline also shows nonlinear trajectories (Wilkosz et al., 2010), one could argue that accounting for nonlinearity is called for when extracting the predictive targets. However, robustly estimating nonlinearity requires more longitudinal measurements per subject and more complex models. In contrast, linear trajectories can robustly be estimated with three measurements, hence, they provide a useful approximation of cognitive decline. Notably, the baseline values of the clinical assessments used to define the slopes have a special role: they are input features and the slopes are defined relative to them. This might result in a bias due to regression to the mean (Barnett et al., 2005), where unusually extreme baseline values (due to noise) might result in unusually extreme slopes (returning to the mean). This issue is relevant as well when defining diagnostic labels where it might result in patients switching between labels due to noise. Future studies should consider more complex models that can better account for these effects. Taken together, quantifying cognitive decline continuously allows for a more nuanced representation of decline and widens the scope of applications. However, while refining the definition of cognitive decline is warranted, it requires more complex analytical approaches and appropriate data.

In this study, we quantified cognitive decline using two clinical assessments (CDR and MMSE), which measure a heterogeneous set of cognitive and everyday life functions. While these clinical assessments have the advantage of being used in practice, they lack the specificity to target single cognitive constructs. Measuring cognitive constructs more homogeneously might potentially improve accuracy, especially if those constructs are strongly linked to specific brain regions or networks. This could be achieved by additionally employing neuropsychological assessments. The multi-target approach outlined in this study is well-suited to including these additional targets.

Beside additional targets, future studies should also consider additional multimodal input data to characterize the brain in greater detail. The present study used data derived from structural and functional MRI (T1w and resting-state fMRI). These might be complemented by information from diffusion-weighted imaging, arterial spin labeling, or positron emission tomography (Rahim et al., 2016). Additionally, the presently used modalities could also be refined and alternative representations could be considered. For instance, different methods for quantifying brain structure (Pipitone et al., 2014) or brain function (Rahim et al., 2019), and adding data on structural asymmetry (Wachinger et al., 2016) or dynamic functional connectivity (Filippi et al., 2019) could provide improved predictive performance. Furthermore, the influence of MR data quality on accuracy should be assessed in future studies. While our past work showed that brain-age prediction from multimodal neuroimaging is robust against in-scanner head motion (Liem et al., 2017), the present study has not assessed the influence of MR data quality on predictive accuracy. Addressing this issue would yield recommendations regarding the required data quality to predict cognitive decline.

The predictive approach could also be expanded to better accommodate high-dimensional data and the messiness of real-world data acquisition. The present study concatenated low-dimensional features across modalities and fed them into one random forest model. Including all features in one model allowed us to consider feature-level interactions across modalities. Alternatively, *prediction stacking* could be used to facilitate the integration of multimodal data (Engemann et al., 2020; Liem et al., 2017; Rahim et al., 2016). While the stacking approach accounts for modality-level interactions it does not consider feature-level interactions across modalities. However, it scales well to high-dimensional data and allows for block-wise missing data, for instance, a missing modality. The present work only included subjects if data from all modalities (non-brain, structural, functional) were available. In clinical practice, this is often not feasible. As we demonstrated previously, stacking can be used to include subjects with missing modalities, which increases the sample size and the scope of application (Engemann et al., 2020).

In practice, the benefit of adding multimodal neuroimaging data to a set of clinical assessments needs to be considered against the additional costs. Its clinical utility also depends on the actionable insight that can be drawn from an earlier prognosis. Of course, this concern is not specific to this study; it applies broadly to almost every effort to incrementally predict clinically meaningful outcomes from brain-based measures. At the moment, no causal treatment of cognitive decline is available. However, an early prognosis might aid intervention studies and be even more helpful once effective treatments are available. Hence, future studies should further exploit the information yielded by the model to focus on subject-specific predictions. In general, predictive models don’t perform equally well in all circumstances. For some subjects or sub-groups a more confident prediction is possible. Recent work demonstrated a higher prediction accuracy in subjects with certain characteristics (e.g., older, female, etc.) (Korolev et al., 2016). This enables increased accuracy by focusing on high-confidence predictions (Tam et al., 2019) and might even suggest a subject-tailored clinical workflow depending on the prediction confidence (Bhagwat et al., 2018). While the present study has not yet investigated these effects, it is well set-up to determine optimal conditions for model performance. The large number of cross-validation splits yields a distribution of predictive performance, not only a point estimate. This will also allow us to assess whether the predictions across sub-groups are driven by the same features.

For a predictive model to be useful in real-world applications, it needs to generalize well to datasets from different sites (Scheinost et al., 2019). While characteristics of our study facilitate generalization, a future study is required to empirically establish the generalization of our models to independent datasets. First, we have aimed to provide full transparency throughout this study to improve reproducibility and generalizability. We used data from a large, publicly available dataset, preprocessed them with well-established open-source tools and inputted them into well-established models. The analysis code is publicly shared and after further developing this approach, trained models will also be shared. Importantly, the analysis was preregistered to avoid overfitting due to analytical flexibility (Carp, 2012; Skocik et al., 2016). Second, the OASIS-3 project is set up heterogeneously regarding the number of sessions, the intervals between sessions, and the participants’ duration in the study. This heterogeneity is expected to provide less opportunity for an algorithm to overfit to dataset-specific idiosyncrasies, resulting in more generalizable models that also perform well in other settings.

While a heterogeneous dataset and open/reproducible approaches certainly improve generalizability, we trained and tested models using only one dataset. Thus, the cross-validated performance in our study provides a biased estimate of the generalizability to independent datasets. This bias might even be modality-specific, in that non-brain features might generalize better than brain imaging features (Bhagwat et al., 2018). Training predictive models on data from multiple sites has been shown to improve generalization (Abraham et al., 2017; Liem et al., 2017; Orban et al., 2018). Hence, future studies should use models trained and tested on data from multiple sites, which requires further suitable longitudinal and publicly available datasets (Varoquaux, 2018). This also provides an opportunity to take preregistration even further. After conducting experiments in an initial dataset, a trained model could be preregistered and applied to an independent dataset that hasn’t yet been analyzed.

## 5 Conclusions

In summary, we have shown that adding structural brain imaging data to non-brain data (such as memory scores or everyday functioning) improves the prediction of future cognitive decline in healthy and pathological aging. Conversely, adding functional connectivity data, as used in the present approach, did not aid the prediction. Importantly, our work has potential for clinical utility by predicting *future* cognitive decline, rather than a *current* diagnosis. Future studies should include additional brain imaging modalities and independent datasets, and should determine the potential of functional connectivity using alternative methodological approaches. Quantifying future decline continuously allows for more nuanced predictions on an individual level. In the future, these continuous measures may facilitate dimensional approaches to pathology (Cuthbert, 2014).

Increased personal and societal costs due to healthy and pathological age-related cognitive decline are one of the most pressing challenges in an aging society. An early and individually fine-grained prognosis of age-related cognitive decline allows for earlier and individually targeted behavioral, cognitive, or pharmacological interventions. Intervening early increases the chances to attenuate or prevent cognitive decline, which will alleviate both personal and societal costs. Importantly, our work targets applications to healthy aging, widening the scope beyond the pathological to the entire aging population.

## Acknowledgments

This work was supported by the URPP “Dynamics of Healthy Aging” at the University of
Zurich. Data were provided by the OASIS-3 project (Principal Investigators: T. Benzinger, D. Marcus, J. Morris; NIH
P50AG00561, P30NS09857781, P01AG026276, P01AG003991, R01AG043434, UL1TR000448, R01EB009352). We thank the participants and organizers of the OASIS-3 project for providing the data.

## 6 Appendix

### 6.1 Supplementary Methods

#### 6.1.1 Details on MRI preprocessing^6^

Results included in this manuscript come from data preprocessed using *fMRIPrep* 1.4.1 (RRID:SCR_016216; Esteban et al., 2018a, 2018b), which is based on *Nipype* 1.2.0 (RRID:SCR_002502; Gorgolewski et al., 2011, 2018).

##### 6.1.1.1 Anatomical data preprocessing

T1-weighted (T1w) images were corrected for intensity non-uniformity (INU) with ‘N4BiasFieldCorrection’ (Tustison et al., 2010), distributed with *ANTs* 2.2.0 (Avants et al., 2008; RRID:SCR_004757; Tustison et al., 2010). The T1w-reference was then skull-stripped with a *Nipype* implementation of the ‘antsBrainExtraction.sh’ workflow (from *ANTs*), using OASIS30ANTs as target template. Brain tissue segmentation of cerebrospinal fluid (CSF), white-matter (WM) and gray-matter (GM) was performed on the brain-extracted T1w using ‘fast’ (*FSL* 5.0.9) (Avants et al., 2008; RRID:SCR_002823; Tustison et al., 2010; Zhang et al., 2001). A T1w-reference map was computed after registration of the T1w images (after INU-correction) using ‘mri_robust_template’ (*FreeSurfer* 6.0.1) (Reuter et al., 2010). Brain surfaces were reconstructed using ‘recon-all’ (*FreeSurfer* 6.0.1) (Dale et al., 1999; RRID:SCR_001847; Reuter et al., 2010), and the brain mask estimated previously was refined with a custom variation of the method to reconcile *ANTs*-derived and *FreeSurfer*-derived segmentations of the cortical gray-matter of Mindboggle (RRID:SCR_002438; Klein et al., 2017). Volume-based spatial normalization to one standard space (MNI152NLin2009cAsym) was performed through nonlinear registration with ‘antsRegistration’ (*ANTs* 2.2.0), using brain-extracted versions of both T1w reference and the T1w template. The following template was selected for spatial normalization: *ICBM 152 Nonlinear Asymmetrical template version 2009c* (Fonov et al., 2009) (RRID:SCR_008796, TemplateFlow ID: MNI152NLin2009cAsym). Where available, T2w-images were included for surface reconstruction.

##### 6.1.1.2 Functional data preprocessing

For each BOLD run, the following preprocessing was performed. First, a reference volume and its skull-stripped version were generated using a custom methodology of *fMRIPrep*. The BOLD reference was then co-registered to the T1w reference using ‘bbregister’ (*FreeSurfer*) which implements boundary-based registration (Greve and Fischl, 2009). Co-registration was configured with nine degrees of freedom to account for distortions remaining in the BOLD reference. Head-motion parameters with respect to the BOLD reference (transformation matrices, and six corresponding rotation and translation parameters) are estimated before any spatiotemporal filtering using ‘mcflirt’ (*FSL* 5.0.9) (Jenkinson et al., 2002). The BOLD time-series (including slice-timing correction) were resampled onto their original, native space by applying a single, composite transform to correct for head-motion and susceptibility distortions. These resampled BOLD time-series will be referred to as *preprocessed BOLD in original space*, or just *preprocessed BOLD*. The BOLD time-series were resampled into standard space, generating a *preprocessed BOLD run in MNI152NLin2009cAsym space*. Several confounding time-series were calculated based on the *preprocessed BOLD*: framewise displacement (FD), DVARS and three region-wise global signals. FD and DVARS are calculated for each functional run, both using their implementations in *Nipype* (following the definitions by (Power et al., 2014). The three global signals are extracted within the CSF, the WM, and the whole-brain masks.

All resamplings can be performed with *a single interpolation step* by composing all the pertinent transformations (i.e. head-motion transform matrices, susceptibility distortion correction when available, and co-registrations to anatomical and output spaces). Gridded (volumetric) resamplings were performed using ‘antsApplyTransforms’ (*ANTs*), configured with Lanczos interpolation to minimize the smoothing effects of other kernels (Lanczos, 1964). Many internal operations of *fMRIPrep* use *Nilearn* 0.5.2 (RRID:SCR_001362; Abraham et al., 2014), mostly within the functional processing workflow. For more details of the pipeline, see the section corresponding to workflows in *fMRIPrep’s* documentation^7^.

#### 6.1.2 Deviation from preregistration

This analysis has been preregistered (Liem et al., 2019). While we have largely followed the plan, the analysis deviates in several minor points:

- Subjects were only considered for the study if they had at least three clinical sessions (not two as preregistered). Extracting the slopes of cognitive decline from two sessions resulted in very noisy slopes. As a result, the sample size is not 849 as preregistered, but 662. As our learning curve experiments demonstrate, the resulting sample size is sufficient in the present context.
- After further discussion, we added two features that have not been preregistered to the non-brain feature set: i) the number of sessions prior to the baseline session (to account for retest effects), and ii) the cognitive diagnosis at baseline (healthy, mild cognitive impairment, dementia). Both features did not show high importance in the permutation importance analysis.
- The preregistered structural features set included 331 features from global, subcortical, and cortical (volume and thickness) markers. After finding that the reduced structGS feature set (35 global and subcortical markers) performs equally well, we conducted the analyses with the more parsimonious structGS modality.
- After further discussion, the preregistered dimensionality reduction approach for the functional connectivity data seemed suboptimal. It averages positive and negative values, which might result in the cancellation of positive and negative connectivity within a network. The updated approach downsampled the connectivity matrix using a PCA.
- In the preregistration, two hyperparameters were planned to be used for tree pruning (max_depth = [5, 10, 20, 40, 50, None], where None leads to fully grown trees and min_samples_leaf = [1, 4, 10]). After further discussion, we decided to remove this double parametrization and only tune max_depth. Additionally, a currently ongoing independent project suggested that lower values of max_depth might be worth investigating in more detail. Hence, we added, max_depth = [3, 7, 15] to the hyperparameter tuning. Since the hyperparameters did not have a large influence on the result and those added values did not render the best performance we deem this change inconsequential.
- The results of the main analysis suggested further validation analyses (brain-age, extreme group classification). Those were not preregistered. However, they are very similar to the main analysis (only the models were adapted to the question: ridge regression for the brain-age analysis; random forest classifier for the extreme group classification with the criterion to measure the quality of an RF-split tuned with [“gini”, “entropy”]).

#### 6.1.3 List of features

**Table S1.** Input features

| Modality | # features modality | Name | Test/Variables | Information | # features |
| --- | --- | --- | --- | --- | --- |
| non-brain | 66 | demographic information | Age, sex, education |  | 3 |
|  |  | clinical assessments | MMSE | Sum score | 1 |
|  |  |  | CDR | 6 items, global and SOB scores | 8 |
|  |  |  | FAQ | 10 items and sum score | 11 |
|  |  |  | NPI-Q | Symptom presence and severity sum scores | 2 |
|  |  |  | GDS | Sum score | 1 |
|  |  | neuropsychology | WMS-R | Logical memory (LOGIMEM, MEMUNITS, MEMTIME)<br>Digit span (DIGIF, DIGIFLEN, DIGIB, DIGIBLEN) | 7 |
|  |  |  | Word fluency | ANIMALS, VEG | 2 |
|  |  |  | TMT | Part A (TRAILA, TRAILARR, TRAILALI), Part B (TRAILB, TRAILBRR, TRAILBLI), TRAILBnorm = TRAILB/TRAILA | 7 |
|  |  |  | WAIS-R | Digit Symbol | 1 |
|  |  |  | BNT | BOSTON | 1 |
| | | APOE | $\epsilon$ 2, $\epsilon$ 3, and $\epsilon$ 4 allele count | | 3 |
|  |  | cognitive diagnosis |  | healthy control, MCI, or dementia | 1 |
|  |  | health | cardio/cerebro-vascular health, diabetes, hypercholesterolemia, smoking, and family history of dementia |  | 17 |
|  |  | N sessions before baseline |  |  | 1 |
| structGS | 35 | subcortical volume | accumbens, amygdala, caudate, hippocampus, pallidum, putamen, thalamus | Volume of left and right | 14 |
|  |  | global measurements | L + R mean cortical thickness; L + R lateral, 3rd, 4th ventricles; L + R total cortical volume; L + R cerebral white matter volume; L + R cerebellar white matter and cortical volume; total subcortical gray matter volume, total gray matter volume, corpus callosum volume (5 parcels) |  | 21 |
| func | 100 | functional connectivity | 300 cortical, cerebellar, and subcortical ROIs | reduced to 100 PCA components | 100 |

### 6.2 Supplementary Results

#### 6.2.1 Predictive targets

**Figure S1.**
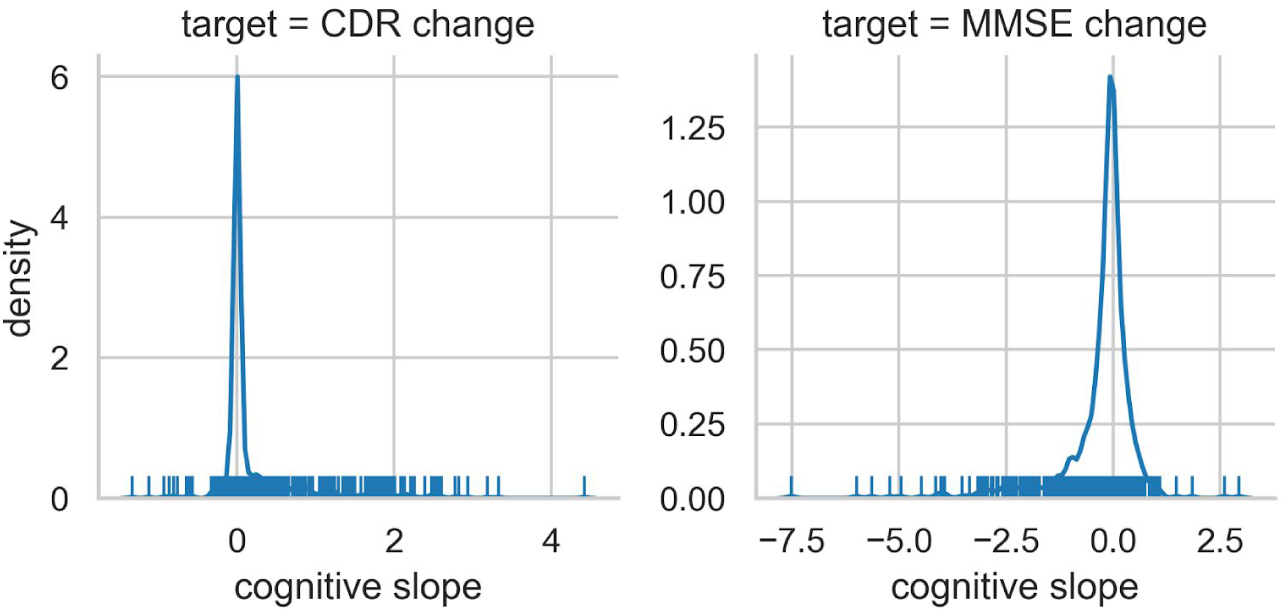
Distribution of predictive targets (cognitive slopes of CDR and MMSE).

#### 6.2.2 Predictive performance (full results)

**Figure S2.**
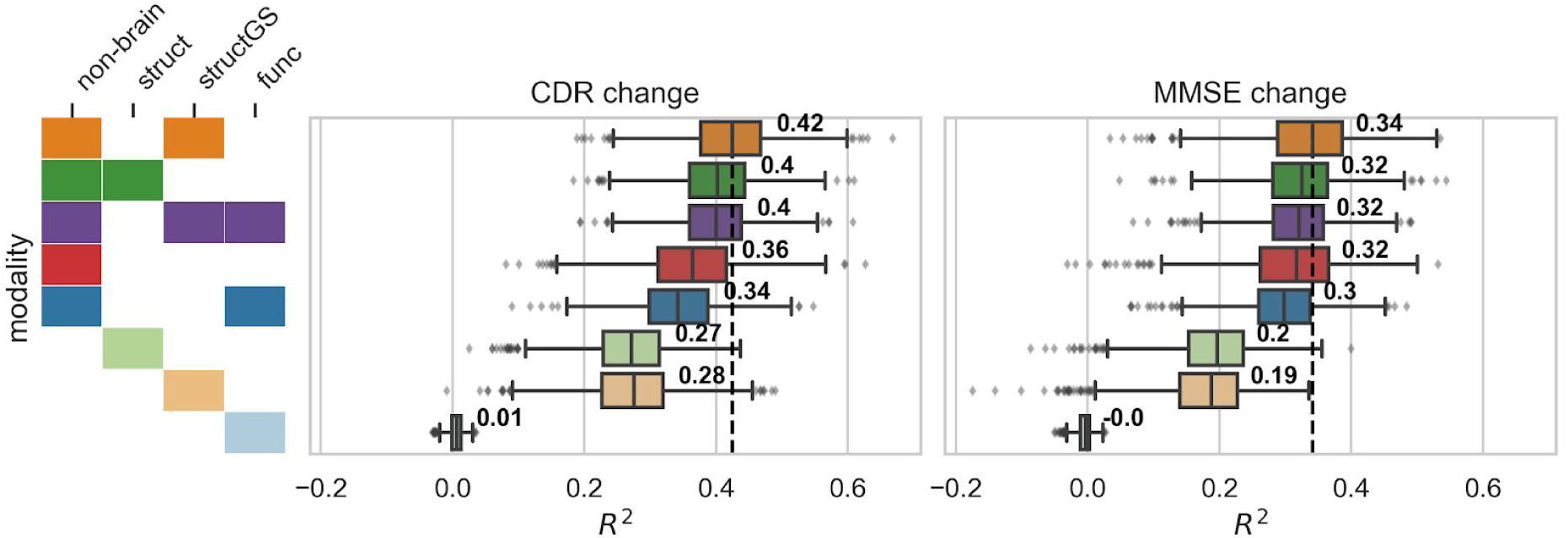
Adding structural data to non-brain data improves prediction of cognitive decline. Test performance (R^2^, coefficient of determination) across splits. Targets: cognitive decline measured via CDR (Clinical Dementia Rating, top) and MMSE (Mini-Mental State Examination, bottom). Input modalities: non-brain, structGS (global and subcortical structural volumes), struct (structGS + cortical volume and thickness), func (functional connectivity). Left panel represents combinations of input modalities (e.g., first line is non-brain + structGS). The number represents the median, the dashed vertical line marks the median of the best-performing combination of modalities (within a target). This figure is an extension of Figure 2 and also includes brain modalities on their own.

**Table S2.** Comparison of test performance vs non-brain predictions (% of splits for which the test prediction R^2^ is outperforming the non-brain prediction; N_splits_ = 1000). Mean and SD difference show the mean and standard deviation of the modality’s performance vs. the non-brain prediction. Best performing model (gray): the CDR prediction from *non-brain + structGS* is outperforming the non-brain prediction in 91% of splits.

| modality / target | outperforming<br>non-brain<br>(% of splits) |  | mean difference |  | SD difference |  |
| --- | --- | --- | --- | --- | --- | --- |
|  | CDR | MMSE | CDR | MMSE | CDR | MMSE |
|  | change | change | change | change | change | change |
| structGS | 19 | 5 | -0.09 | -0.13 | 0.1 | 0.09 |
| struct | 17 | 7 | -0.09 | -0.12 | 0.1 | 0.09 |
| func | 0 | 0 | -0.36 | -0.31 | 0.08 | 0.08 |
| non-brain + structGS | 92 | 78 | 0.06 | 0.02 | 0.04 | 0.04 |
| non-brain + struct | 75 | 59 | 0.04 | 0.01 | 0.06 | 0.05 |
| non-brain + func | 25 | 31 | -0.02 | -0.01 | 0.03 | 0.03 |
| non-brain + structGS +<br>func | 78 | 57 | 0.04 | 0.01 | 0.05 | 0.04 |

**Table S3.** Comparison of test performance vs null-models (% of splits for which the test prediction R^2^ is outperforming the null-model prediction). Best performing model (gray): the CDR prediction from *non-brain + structGS* is outperforming the non-brain prediction in 100% of splits.

| modality / target | outperforming<br>null-models<br>(% of splits) |  | median difference |  | SD difference |  |
| --- | --- | --- | --- | --- | --- | --- |
|  | CDR | MMSE | CDR | MMSE | CDR | MMSE |
|  | change | change | change | change | change | change |
| non-brain | 100 | 100 | 0.42 | 0.36 | 0.09 | 0.09 |
| structGS | 100 | 99 | 0.31 | 0.21 | 0.08 | 0.07 |
| struct | 100 | 100 | 0.34 | 0.24 | 0.08 | 0.07 |
| func | 91 | 73 | 0.01 | 0.01 | 0.01 | 0.01 |
| non-brain + structGS | 100 | 100 | 0.48 | 0.39 | 0.08 | 0.09 |
| non-brain + struct | 100 | 100 | 0.47 | 0.38 | 0.09 | 0.09 |
| non-brain + func | 100 | 100 | 0.42 | 0.35 | 0.08 | 0.07 |
| non-brain + structGS +<br>func | 100 | 100 | 0.47 | 0.37 | 0.08 | 0.08 |

**Figure S3.**
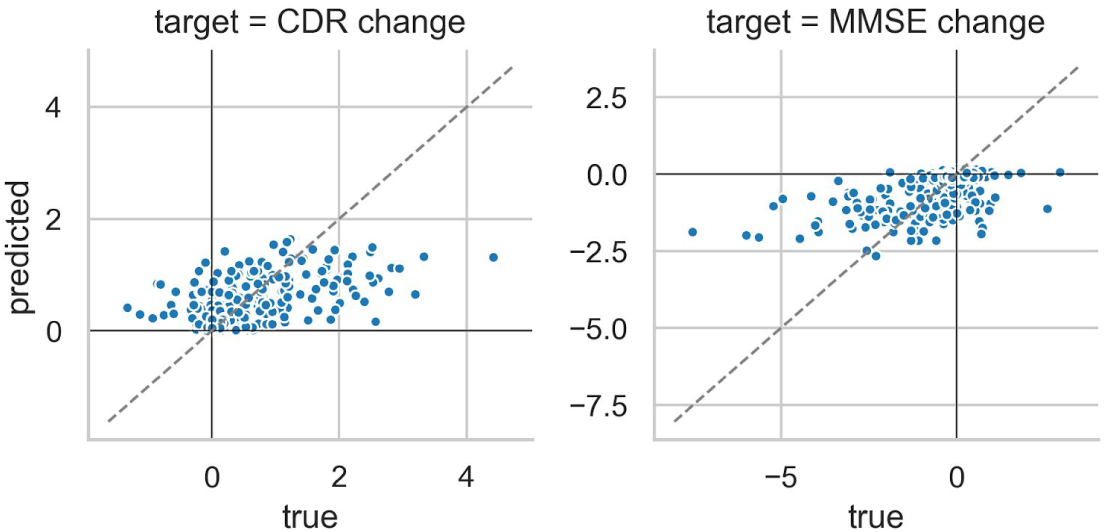
Scatter plot of true vs predicted trajectories of cognitive change for the *non-brain + structGS* model. Mean prediction across splits. CDR: positive values = cognitive decline, MMSE: negative values = cognitive decline.

#### 6.2.3 Hyperparameter tuning of the random forest regression model

**Figure S4.**
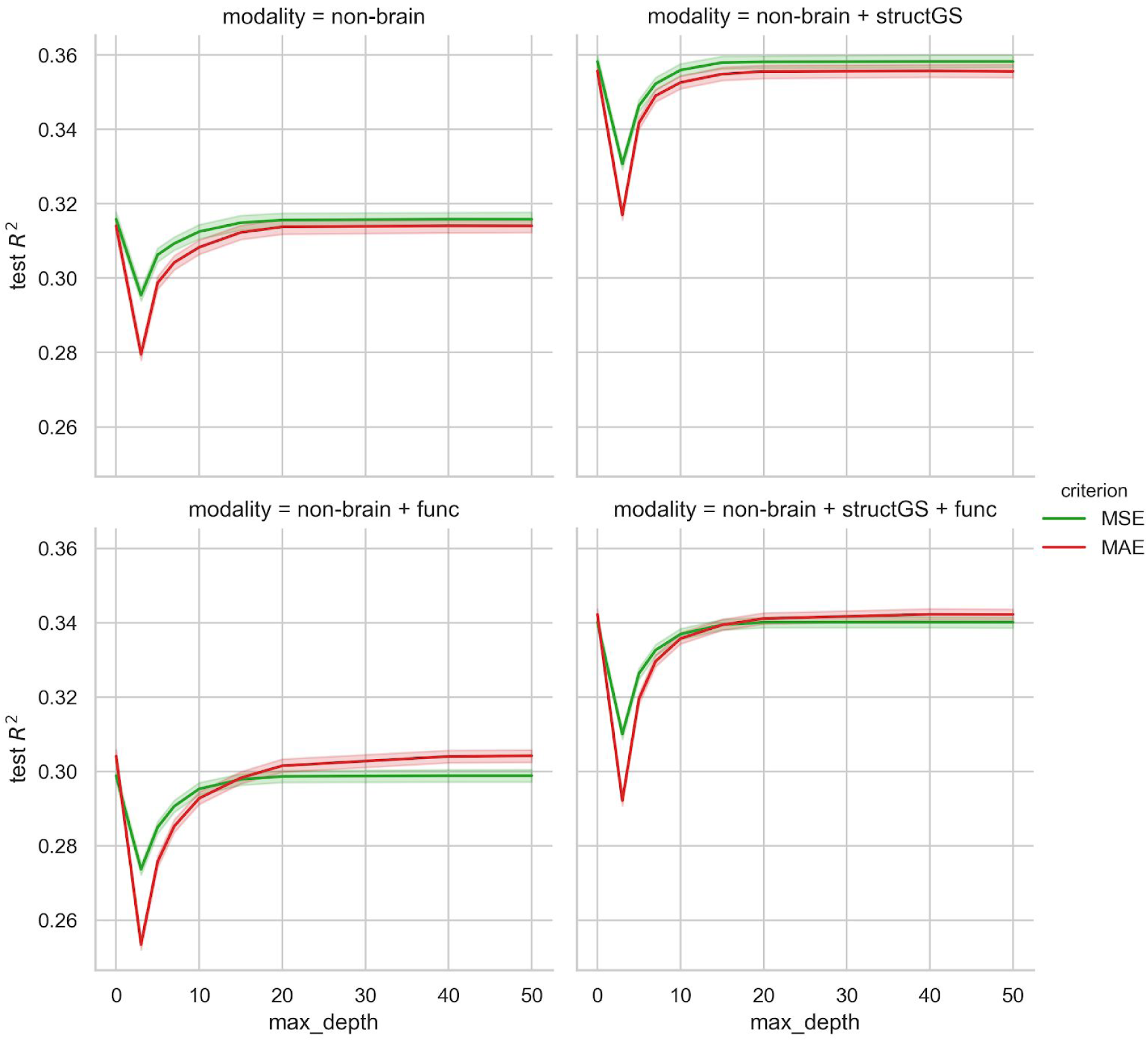
Tuning curves of the random forest regression hyperparameters max tree depth (0 represents fully grown trees) and criterion (MSE: mean squared error, MAE: mean absolute error). Note that the y-axis is trimmed and the results vary in a rather narrow range.

#### 6.2.4 Learning curves demonstrate sufficient sample size

**Figure S5.**
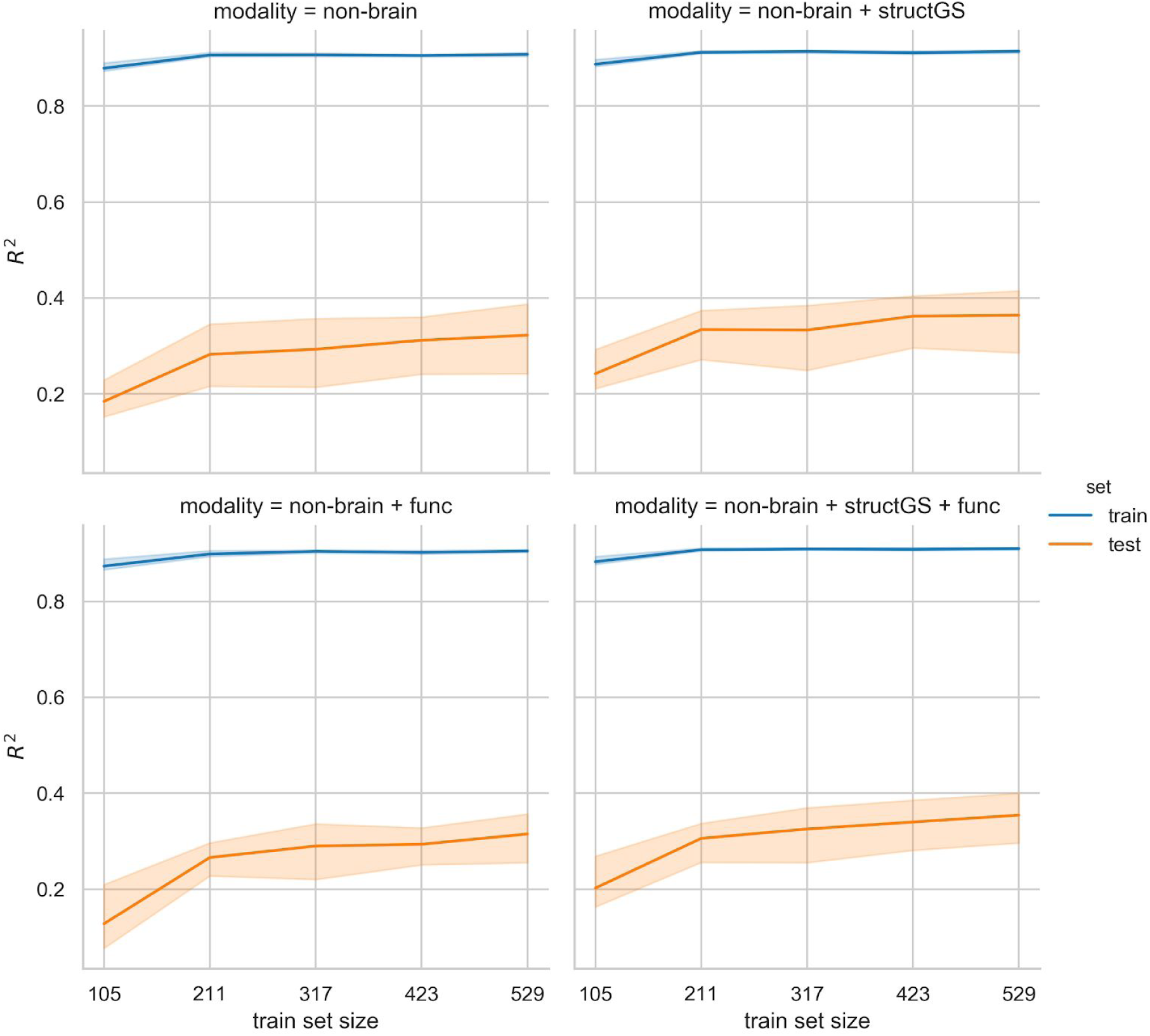
Learning curve plotting the models’ performance across an increasing training sample size.

#### 6.2.5 Age prediction

**Figure S6.**
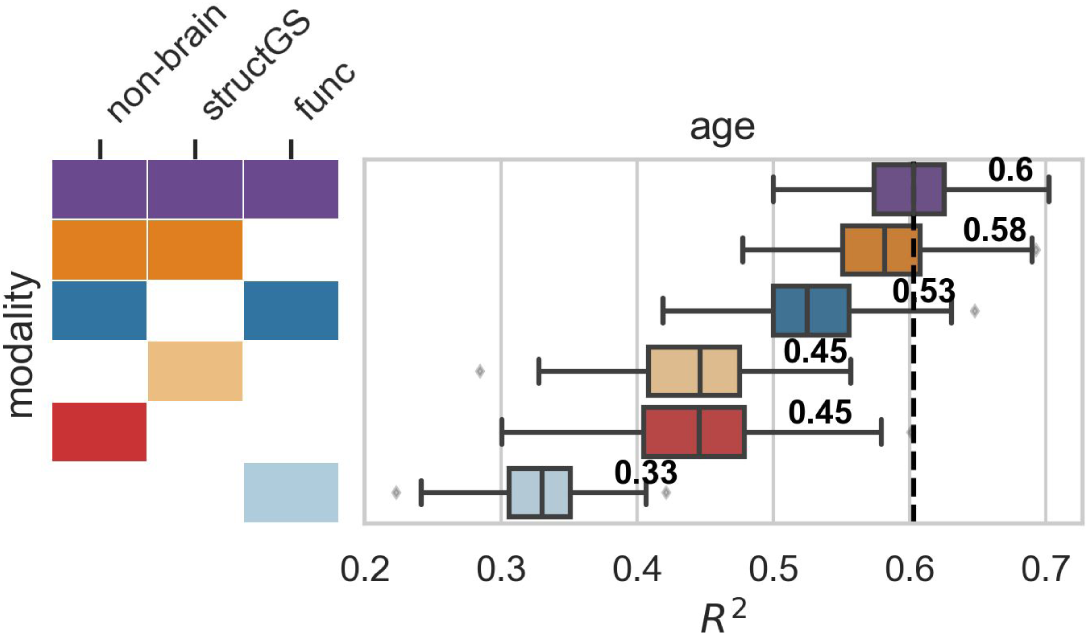
Predictive performance in brain-age prediction. Input modalities: non-brain, structGS (global and subcortical structural volumes), func (functional connectivity). The number represents the median, the dashed vertical line marks the median of the best-performing combination of modalities. This figure is an extension of Figure 4 and also includes brain modalities on their own.

#### 6.2.6 Extreme group prediction

**Figure S7.**
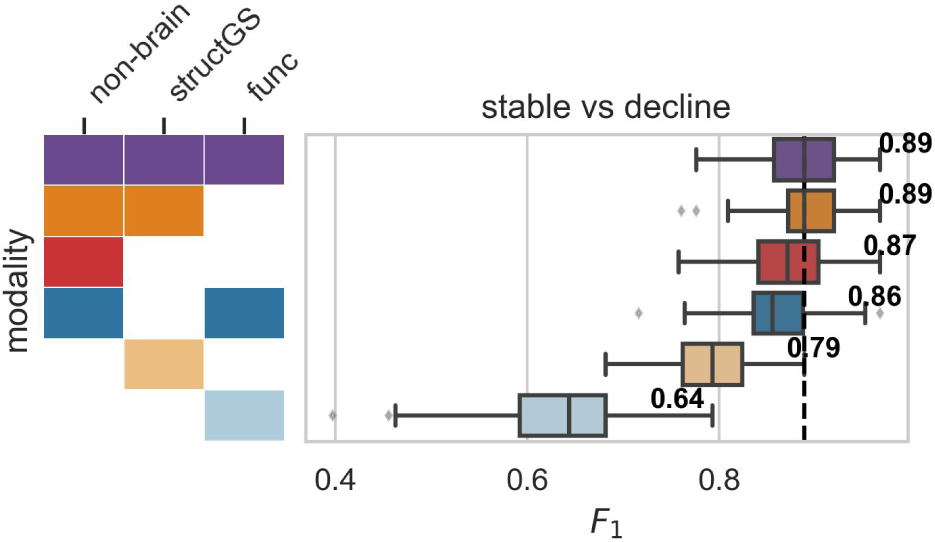
Predictive performance in predicting extreme groups of cognitive decline (stable vs decline; F1-score: harmonic mean of the precision and recall). Input modalities: non-brain, structGS (global and subcortical structural volumes), func (functional connectivity). The number represents the median, the dashed vertical line marks the median of the best-performing combination of modalities.

## Footnotes

1 The selected subjects and session can be found here: https://github.com/fliem/cpr/tree/0.1.2/info

2 https://github.com/NrgXnat/oasis-scripts

3 https://central.xnat.org/

4 http://github.com/fliem/cpr

5 https://hub.docker.com/r/fliem/cpr

6 The description in this section was automatically created by *fMRIPrep* and adapted where needed.

7 https://fmriprep.readthedocs.io/en/latest/workflows.html

## References

Abraham A, Milham MP, Di Martino A, Craddock RC, Samaras D, Thirion B, Varoquaux G. 2017. Deriving reproducible biomarkers from multi-site resting-state data: An Autism-based example. Neuroimage 147:736–745. doi: 10.1016/j.neuroimage.2016.10.045

Abraham A, Pedregosa F, Eickenberg M, Gervais P, Mueller A, Kossaifi J, Gramfort A, Thirion B, Varoquaux G. 2014. Machine learning for neuroimaging with scikit-learn. Front Neuroinform 8. doi: 10.3389/fninf.2014.00014

Avants BB, Epstein CL, Grossman M, Gee JC. 2008. Symmetric diffeomorphic image registration with cross-correlation: Evaluating automated labeling of elderly and neurodegenerative brain. Med Image Anal 12: 26–41. doi: 10.1016/j.media.2007.06.004

Barnett AG, van der Pols JC, Dobson AJ. 2005. Regression to the mean: what it is and how to deal with it. Int J Epidemiol 34:215–220. doi: 10.1093/ije/dyh299

Bengio Y, Grandvalet Y. 2004. No Unbiased Estimator of the Variance of K-Fold Cross-Validation. J Mach Learn Res 5:1089–1105.

Bhagwat N, Viviano JD, Voineskos AN, Chakravarty MM, Alzheimer’s Disease Neuroimaging Initiative. 2018. Modeling and prediction of clinical symptom trajectories in Alzheimer’s disease using longitudinal data. PLoS Comput Biol 14:e1006376. doi: 10.1371/journal.pcbi.1006376

Borod JC, Goodglass H, Kaplan E. 1980. Normative data on the boston diagnostic aphasia examination, parietal lobe battery, and the boston naming Test. Journal of Clinical Neuropsychology. doi: 10.1080/01688638008403793

Breiman L. 2001. 10.1023/A:1010933404324. Machine Learning. doi: 10.1023/A:1010933404324

Buck SF. 1960. A Method of Estimation of Missing Values in Multivariate Data Suitable for use with an Electronic Computer. J R Stat Soc Series B Stat Methodol 22:302–306.

Carp J. 2012. On the plurality of (methodological) worlds: estimating the analytic flexibility of FMRI experiments. Front Neurosci 6:149. doi: 10.3389/fnins.2012.00149

Ciric R, Wolf DH, Power JD, Roalf DR, Baum GL, Ruparel K, Shinohara RT, Elliott MA, Eickhoff SB, Davatzikos C, Gur RC, Gur RE, Bassett DS, Satterthwaite TD. 2017. Benchmarking of participant-level confound regression strategies for the control of motion artifact in studies of functional connectivity. Neuroimage 154:174–187. doi: 10.1016/j.neuroimage.2017.03.020

Cole JH, Franke K. 2017. Predicting Age Using Neuroimaging: Innovative Brain Ageing Biomarkers. Trends Neurosci 40:681–690. doi: 10.1016/j.tins.2017.10.001

Cuthbert BN. 2014. The RDoC framework: facilitating transition from ICD/DSM to dimensional approaches that integrate neuroscience and psychopathology. World Psychiatry 13:28–35. doi: 10.1002/wps.20087

Dale AM, Fischl B, Sereno MI. 1999. Cortical Surface-Based Analysis: I. Segmentation and Surface Reconstruction. Neuroimage 9:179–194. doi: 10.1006/nimg.1998.0395

Dansereau C, Tam A, Badhwar A, Urchs S, Orban P, Rosa-Neto P, Bellec P. 2017. A brain signature highly predictive of future progression to Alzheimer’s dementia.

Davatzikos C. 2019. Machine learning in neuroimaging: Progress and challenges. Neuroimage 197:652–656. doi: 10.1016/j.neuroimage.2018.10.003

Davatzikos C, Bhatt P, Shaw LM, Batmanghelich KN, Trojanowski JQ. 2011. Prediction of MCI to AD conversion, via MRI, CSF biomarkers, and pattern classification. Neurobiol Aging 32:2322.e19–27. doi: 10.1016/j.neurobiolaging.2010.05.023

Desikan RS, Ségonne F, Fischl B, Quinn BT, Dickerson BC, Blacker D, Buckner RL, Dale AM, Maguire RP, Hyman BT, Albert MS, Killiany RJ. 2006. An automated labeling system for subdividing the human cerebral cortex on MRI scans into gyral based regions of interest. Neuroimage 31:968–980. doi: 10.1016/j.neuroimage.2006.01.021

Dubois J, Galdi P, Han Y, Paul LK, Adolphs R. 2018. Resting-State Functional Brain Connectivity Best Predicts the Personality Dimension of Openness to Experience. Personality Neuroscience. doi: 10.1017/pen.2018.8

Elwood RW. 1991. The Wechsler Memory Scale-Revised: psychometric characteristics and clinical application. Neuropsychol Rev 2:179–201. doi: 10.1007/bf01109053

Engemann DA, Kozynets O, Sabbagh D, Lemaître G, Varoquaux G, Liem F, Gramfort A. 2020. Combining magnetoencephalography with magnetic resonance imaging enhances learning of surrogate-biomarkers. Elife 9. doi: 10.7554/eLife.54055

Eskildsen SF, Coupé P, Fonov VS, Pruessner JC, Collins DL, Alzheimer’s Disease Neuroimaging Initiative. 2015. Structural imaging biomarkers of Alzheimer’s disease: predicting disease progression. Neurobiol Aging 36 Suppl 1: S23–31. doi: 10.1016/j.neurobiolaging.2014.04.034

Esteban O, Blair R, Markiewicz CJ, Berleant SL, Moodie C, Ma F, Isik AI, Erramuzpe A, Kent M James D andGoncalves, DuPre E, Sitek KR, Gomez DEP, Lurie DJ, Ye Z, Poldrack RA, Gorgolewski KJ. 2018a. fMRIPrep. Softw Pract Exp. doi: 10.5281/zenodo.852659

Esteban O, Markiewicz C, Blair RW, Moodie C, Isik AI, Erramuzpe Aliaga A, Kent J, Goncalves M, DuPre E, Snyder M, Oya H, Ghosh S, Wright J, Durnez J, Poldrack R, Gorgolewski KJ. 2018b. fMRIPrep: a robust preprocessing pipeline for functional MRI. Nat Methods. doi: 10.1038/s41592-018-0235-4

Filippi M, Spinelli EG, Cividini C, Agosta F. 2019. Resting State Dynamic Functional Connectivity in Neurodegenerative Conditions: A Review of Magnetic Resonance Imaging Findings. Front Neurosci 13:657. doi: 10.3389/fnins.2019.00657

Fischl B. 2012. FreeSurfer. NeuroImage. doi: 10.1016/j.neuroimage.2012.01.021

Folstein MF, Folstein SE, McHugh PR. 1975. “Mini-mental state”. A practical method for grading the cognitive state of patients for the clinician. J Psychiatr Res 12:189–198.

Fonov VS, Evans AC, McKinstry RC, Almli CR, Collins DL. 2009. Unbiased nonlinear average age-appropriate brain templates from birth to adulthood. Neuroimage 47, Supplement 1: S102. doi: 10.1016/S1053-8119(09)70884-5

Franzen MD. 2000. The Wechsler Adult Intelligence Scale-Revised and Wechsler Adult Intelligence Scale-III. Reliability and Validity in Neuropsychological Assessment. doi: 10.1007/978-1-4757-3224-5_6

Gaser C, Franke K, Klöppel S, Koutsouleris N, Sauer H, Alzheimer’s Disease Neuroimaging Initiative. 2013. BrainAGE in Mild Cognitive Impaired Patients: Predicting the Conversion to Alzheimer’s Disease. PLoS One 8:e67346. doi: 10.1371/journal.pone.0067346

Gorgolewski K, Burns CD, Madison C, Clark D, Halchenko YO, Waskom ML, Ghosh S. 2011. Nipype: a flexible, lightweight and extensible neuroimaging data processing framework in Python. Front Neuroinform 5:13. doi: 10.3389/fninf.2011.00013

Gorgolewski KJ, Auer T, Calhoun VD, Craddock RC, Das S, Duff EP, Flandin G, Ghosh SS, Glatard T, Halchenko YO, Handwerker DA, Hanke M, Keator D, Li X, Michael Z, Maumet C, Nichols BN, Nichols TE, Pellman J, Poline J-B, Rokem A, Schaefer G, Sochat V, Triplett W, Turner JA, Varoquaux G, Poldrack RA. 2016. The brain imaging data structure, a format for organizing and describing outputs of neuroimaging experiments. Sci Data 3:160044. doi: 10.1038/sdata.2016.44

Gorgolewski KJ, Esteban O, Markiewicz CJ, Ziegler E, Ellis DG, Notter MP, Jarecka D, Johnson H, Burns C, Manhães-Savio A, Hamalainen C, Yvernault B, Salo T, Jordan K, Goncalves M, Waskom M, Clark D, Wong J, Loney F, Modat M, Dewey BE, Madison C, Visconti di Oleggio Castello M, Clark MG, Dayan M, Clark D, Keshavan A, Pinsard B, Gramfort A, Berleant S, Nielson DM, Bougacha S, Varoquaux G, Cipollini B, Markello R, Rokem A, Moloney B, Halchenko YO, Wassermann D, Hanke M, Horea C, Kaczmarzyk J, de Hollander G, DuPre E, Gillman A, Mordom D, Buchanan C, Tungaraza R, Pauli WM, Iqbal S, Sikka S, Mancini M, Schwartz Y, Malone IB, Dubois M, Frohlich C, Welch D, Forbes J, Kent J, Watanabe A, Cumba C, Huntenburg JM, Kastman E, Nichols BN, Eshaghi A, Ginsburg D, Schaefer A, Acland B, Giavasis S, Kleesiek J, Erickson D, Küttner R, Haselgrove C, Correa C, Ghayoor A, Liem F, Millman J, Haehn D, Lai J, Zhou D, Blair R, Glatard T, Renfro M, Liu S, Kahn AE, Pérez-García F, Triplett W, Lampe L, Stadler J, Kong X-Z, Hallquist M, Chetverikov A, Salvatore J, Park A, Poldrack R, Craddock RC, Inati S, Hinds O, Cooper G, Perkins LN, Marina A, Mattfeld A, Noel M, Snoek L, Matsubara K, Cheung B, Rothmei S, Urchs S, Durnez J, Mertz F, Geisler D, Floren A, Gerhard S, Sharp P, Molina-Romero M, Weinstein A, Broderick W, Saase V, Andberg SK, Harms R, Schlamp K, Arias J, Papadopoulos Orfanos D, Tarbert C, Tambini A, De La Vega A, Nickson T, Brett M, Falkiewicz M, Podranski K, Linkersdörfer J, Flandin G, Ort E, Shachnev D, McNamee D, Davison A, Varada J, Schwabacher I, Pellman J, Perez-Guevara M, Khanuja R, Pannetier N, McDermottroe C, Ghosh S. 2018. Nipype. Softw Pract Exp. doi: 10.5281/zenodo.596855

Greve DN, Fischl B. 2009. Accurate and robust brain image alignment using boundary-based registration. Neuroimage 48:63–72. doi: 10.1016/j.neuroimage.2009.06.060

Heller LJ, Skinner CS, Tomiyama AJ, Epel ES, Hall PA, Allan J, LaCaille L, Randall AK, Bodenmann G, Li-Tsang CWP, Sinclair K, Creek J, Baumann LC, Karel A, Andersson G, Hanewinkel R, Morgenstern M, Puska P, Bucks RS, Carroll J, Gidron Y, Rosenberg L, Delamater AM, Gidron Y, Spring B, Coons MJ, Duncan J, Sularz A, Deary J, Prochaska JO, Spiers M, Reid EW, Miller R, Kirschbaum C, Barrett C, Rohleder N, Wessel J, Meneghini L, Pulgaron ER, Delamater AM, Suzuki S-I, Kunisato Y, Denollet J. 2013. Trail-Making Test In: Gellman MD, Turner JR, editors. Encyclopedia of Behavioral Medicine. New York, NY: Springer New York. pp. 1986–1987. doi: 10.1007/978-1-4419-1005-9_1538

Hojjati SH, Ebrahimzadeh A, Khazaee A, Babajani-Feremi A, Alzheimer’s Disease Neuroimaging Initiative. 2018. Predicting conversion from MCI to AD by integrating rs-fMRI and structural MRI. Comput Biol Med 102:30–39. doi: 10.1016/j.compbiomed.2018.09.004

Jenkinson M, Bannister P, Brady M, Smith S. 2002. Improved Optimization for the Robust and Accurate Linear Registration and Motion Correction of Brain Images. Neuroimage 17:825–841. doi: 10.1006/nimg.2002.1132

Jette AM, Davies AR, Cleary PD, Calkins DR, Rubenstein LV, Fink A, Kosecoff J, Young RT, Brook RH, Delbanco TL. 1986. The Functional Status Questionnaire: reliability and validity when used in primary care. J Gen Intern Med 1:143–149.

Josse J, Prost N, Scornet E, Varoquaux G. 2019. On the consistency of supervised learning with missing values.

Kaufer DI, Cummings JL, Ketchel P, Smith V, MacMillan A, Shelley T, Lopez OL, DeKosky ST. 2000. Validation of the NPI-Q, a brief clinical form of the Neuropsychiatric Inventory. J Neuropsychiatry Clin Neurosci 12:233–239. doi: 10.1176/jnp.12.2.233

Klein A, Ghosh SS, Bao FS, Giard J, Häme Y, Stavsky E, Lee N, Rossa B, Reuter M, Neto EC, Keshavan A. 2017. Mindboggling morphometry of human brains. PLoS Comput Biol 13:e1005350. doi: 10.1371/journal.pcbi.1005350

Korolev IO, Symonds LL, Bozoki AC, Alzheimer’s Disease Neuroimaging Initiative. 2016. Predicting Progression from Mild Cognitive Impairment to Alzheimer’s Dementia Using Clinical, MRI, and Plasma Biomarkers via Probabilistic Pattern Classification. PLoS One 11:e0138866. doi: 10.1371/journal.pone.0138866

LaMontagne PJ, Benzinger TLS, Morris JC, Keefe S, Hornbeck R, Xiong C, Grant E, Hassenstab J, Moulder K, Vlassenko A, Raichle ME, Cruchaga C, Marcus D. 2019. OASIS-3: Longitudinal Neuroimaging, Clinical, and Cognitive Dataset for Normal Aging and Alzheimer Disease. Radiology and Imaging.

Lanczos C. 1964. Evaluation of Noisy Data. Journal of the Society for Industrial and Applied Mathematics Series B Numerical Analysis 1:76–85. doi: 10.1137/0701007

Liem F. 2020. fliem/cpr 0.1.1. doi: 10.5281/zenodo.3726641

Liem F, Bellec P, Craddock C, Kamalaker Dadi, Damoiseaux JS, Margulies DS, Steele CJ, Varoquaux G, Yarkoni T. 2019. Predicting future cognitive change from multiple data sources (pilot study).

Liem F, Geerligs L, Damoiseaux JS, Margulies DS. 2020. Functional Connectivity in Aging. doi: 10.31234/osf.io/whsud

Liem F, Varoquaux G, Kynast J, Beyer F, Kharabian Masouleh S, Huntenburg JM, Lampe L, Rahim M, Abraham A, Craddock RC, Riedel-Heller S, Luck T, Loeffler M, Schroeter ML, Witte AV, Villringer A, Margulies DS. 2017. Predicting brain-age from multimodal imaging data captures cognitive impairment. Neuroimage 148:179–188. doi: 10.1016/j.neuroimage.2016.11.005

Morris JC. 1993. The Clinical Dementia Rating (CDR): current version and scoring rules. Neurology 43:2412–2414.

Morris JC, Weintraub S, Chui HC, Cummings J, Decarli C, Ferris S, Foster NL, Galasko D, Graff-Radford N, Peskind ER, Beekly D, Ramos EM, Kukull WA. 2006. The Uniform Data Set (UDS): clinical and cognitive variables and descriptive data from Alzheimer Disease Centers. Alzheimer Dis Assoc Disord 20:210–216. doi: 10.1097/01.wad.0000213865.09806.92

Orban P, Dansereau C, Desbois L, Mongeau-Pérusse V, Giguère C-É, Nguyen H, Mendrek A, Stip E, Bellec P. 2018. Multisite generalizability of schizophrenia diagnosis classification based on functional brain connectivity. Schizophr Res 192:167–171. doi: 10.1016/j.schres.2017.05.027

Oschwald J, Guye S, Liem F, Rast P, Willis S, Röcke C, Jäncke L, Martin M, Mérillat S. 2019. Brain structure and cognitive ability in healthy aging: a review on longitudinal correlated change. Rev Neurosci 31:1–57. doi: 10.1515/revneuro-2018-0096

Pedregosa F, Varoquaux G, Gramfort A, Michel V, Thirion B, Grisel O, Blondel M, Prettenhofer P, Weiss R, Dubourg V, Vanderplas J, Passos A, Cournapeau D, Brucher M, Perrot M, Duchesnay É. 2011. Scikit-learn: Machine Learning in Python. J Mach Learn Res 12:2825–2830.

Pipitone J, Park MTM, Winterburn J, Lett TA, Lerch JP, Pruessner JC, Lepage M, Voineskos AN, Chakravarty MM, Alzheimer’s Disease Neuroimaging Initiative. 2014. Multi-atlas segmentation of the whole hippocampus and subfields using multiple automatically generated templates. Neuroimage 101:494–512. doi: 10.1016/j.neuroimage.2014.04.054

Power JD, Mitra A, Laumann TO, Snyder AZ, Schlaggar BL, Petersen SE. 2014. Methods to detect, characterize, and remove motion artifact in resting state fMRI. Neuroimage 84:320–341. doi: 10.1016/j.neuroimage.2013.08.048

Rahim M, Thirion B, Bzdok D, Buvat I, Varoquaux G. 2017. Joint prediction of multiple scores captures better individual traits from brain images. Neuroimage 158:145–154. doi: 10.1016/j.neuroimage.2017.06.072

Rahim M, Thirion B, Comtat C, Varoquaux G, Alzheimer’s Disease Neuroimaging Initiative. 2016. Transmodal Learning of Functional Networks for Alzheimer’s Disease Prediction. IEEE J Sel Top Signal Process 10:120–1213. doi: 10.1109/JSTSP.2016.2600400

Rahim M, Thirion B, Varoquaux G. 2019. Population shrinkage of covariance (PoSCE) for better individual brain functional-connectivity estimation. Med Image Anal 54:138–148. doi: 10.1016/j.media.2019.03.001

Rathore S, Habes M, Iftikhar MA, Shacklett A, Davatzikos C. 2017. A review on neuroimaging-based classification studies and associated feature extraction methods for Alzheimer’s disease and its prodromal stages. Neuroimage 155:530–548. doi: 10.1016/j.neuroimage.2017.03.057

Reuter M, Rosas HD, Fischl B. 2010. Highly accurate inverse consistent registration: A robust approach. Neuroimage 53:1181–1196. doi: 10.1016/j.neuroimage.2010.07.020

Scheinost D, Noble S, Horien C, Greene AS, Lake EM, Salehi M, Gao S, Shen X, O’Connor D, Barron DS, Yip SW, Rosenberg MD, Constable RT. 2019. Ten simple rules for predictive modeling of individual differences in neuroimaging. Neuroimage 193:35–45. doi: 10.1016/j.neuroimage.2019.02.057

Seabold S, Perktold J. 2010. Statsmodels: Econometric and statistical modeling with pythonProceedings of the 9th Python in Science Conference. Scipy. p. 61.

Seitzman BA, Gratton C, Marek S, Raut RV, Dosenbach NUF, Schlaggar BL, Petersen SE, Greene DJ. 2018. A set of functionally-defined brain regions with improved representation of the subcortex and cerebellum. Neuroscience.

Skocik M, Collins J, Callahan-Flintoft C, Bowman H, Wyble B. 2016. I TRIED A BUNCH OF THINGS: THE DANGERS OF UNEXPECTED OVERFITTING IN CLASSIFICATION. Neuroscience.

Tam A, Dansereau C, Iturria-Medina Y, Urchs S, Orban P, Sharmarke H, Breitner J, Bellec P, Alzheimer’s Disease Neuroimaging Initiative. 2019. A highly predictive signature of cognition and brain atrophy for progression to Alzheimer’s dementia. Gigascience 8. doi: 10.1093/gigascience/giz055

Tustison NJ, Avants BB, Cook PA, Zheng Y, Egan A, Yushkevich PA, Gee JC. 2010. N4ITK: Improved N3 Bias Correction. IEEE Trans Med Imaging 29:1310–1320. doi: 10.1109/TMI.2010.2046908

Varoquaux G. 2018. Cross-validation failure: Small sample sizes lead to large error bars. Neuroimage 180:68–77. doi: 10.1016/j.neuroimage.2017.06.061

Vogel JW, Vachon-Presseau E, Pichet Binette A, Tam A, Orban P, La Joie R, Savard M, Picard C, Poirier J, Bellec P, Breitner JCS, Villeneuve S, Alzheimer’s Disease Neuroimaging Initiative* and the PREVENT-AD Research Group. 2018. Brain properties predict proximity to symptom onset in sporadic Alzheimer’s disease. Brain 141:1871–1883. doi: 10.1093/brain/awy093

Wachinger C, Salat DH, Weiner M, Reuter M, Alzheimer’s Disease Neuroimaging Initiative. 2016. Whole-brain analysis reveals increased neuroanatomical asymmetries in dementia for hippocampus and amygdala. Brain 139:3253–3266. doi: 10.1093/brain/aww243

Weintraub S, Salmon D, Mercaldo N, Ferris S, Graff-Radford NR, Chui H, Cummings J, DeCarli C, Foster NL, Galasko D, Peskind E, Dietrich W, Beekly DL, Kukull WA, Morris JC. 2009. The Alzheimer’s Disease Centers’ Uniform Data Set (UDS): the neuropsychologic test battery. Alzheimer Dis Assoc Disord 23:91–101. doi: 10.1097/WAD.0b013e318191c7dd

Wilkosz PA, Seltman HJ, Devlin B, Weamer EA, Lopez OL, DeKosky ST, Sweet RA. 2010. Trajectories of cognitive decline in Alzheimer’s disease. Int Psychogeriatr 22:281–290. doi: 10.1017/S1041610209991001

Woo C-W, Chang LJ, Lindquist MA, Wager TD. 2017. Building better biomarkers: brain models in translational neuroimaging. Nat Neurosci 20:365–377. doi: 10.1038/nn.4478

Yesavage JA, Brink TL, Rose TL, Lum O, Huang V, Adey M, Leirer VO. 1982. Development and validation of a geriatric depression screening scale: a preliminary report. J Psychiatr Res 17:37–49.

Zhang Y, Brady M, Smith S. 2001. Segmentation of brain MR images through a hidden Markov random field model and the expectation-maximization algorithm. IEEE Trans Med Imaging 20:45–57. doi: 10.1109/42.906424

